# dFCExpert: Learning Dynamic Functional Connectivity Patterns with Modularity and State Experts

**DOI:** 10.1101/2024.12.20.629773

**Authors:** Tingting Chen, Hongming Li, Hao Zheng, Yong Fan

**Author notes:** / /).

## Abstract

Characterizing brain dynamic functional connectivity (dFC) patterns from functional Magnetic Resonance Imaging (fMRI) data is of paramount importance in imaging neuroscience and medicine. Recently, many graph neural network (GNN) models, combined with transformers or recurrent neural networks (RNNs), have shown great potential for modeling the dFC patterns. However, these methods face challenges in effectively characterizing the modularity organization of brain networks and capturing varying dFC state patterns. To address these limitations, we propose dFCExpert, a novel method designed to learn robust representations of dFC patterns in fMRI data with modularity experts and state experts. Specifically, the modularity experts optimize multiple experts to characterize the brain modularity organization during graph feature learning process by combining GNN and mixture of experts (MoE), with each expert focusing on brain nodes within the same functional network module. The state experts aggregate temporal dFC features into a set of distinctive connectivity states using a soft prototype clustering method, providing insight into how these states support diverse brain functions and how they vary across brain conditions. Experiments on three large-scale fMRI datasets have demonstrated the superiority of our method over existing alternatives. The learned dFC representations not only enhance interpretability but also hold promise for advancing our understanding of brain function across a range of conditions, including development, sex difference, and Autism Spectrum Disorder. Our code is publicly available at MLDataAnalytics/dFCExpert^1^.

## I. Introduction

The human brain is a dynamic network system that generates complex spatiotemporal dynamics of brain activity. Analyzing such dynamics holds promise to provide insights into the brain’s functional organization and its relationship with human cognition, behaviors, and brain disorders [1]–[5]. Among many brain imaging techniques, functional Magnetic Resonance Imaging (fMRI) is a particularly powerful tool that models the spatiotemporal patterns of brain activity by measuring fluctuations in blood-oxygen level-dependent (BOLD) signals [6]. Since brain activity often exhibits strong spatial correlations, BOLD signals can be parcellated into a set of pre-defined brain regions of interest (ROIs). The pairwise correlations between those ROIs define functional connectivity (FC), which has emerged as a key tool for understanding brain function.

Based on the FC, the brain can be modeled as a functional network with graph theory approaches, where brain ROIs serve as network nodes and the FC strengths between them act as edges. Leveraging this graph-structured nature of the brain, many fMRI data analytic methods have employed graph neural networks (GNNs) to learn brain network representations, enabling tasks such as decoding human traits or diagnosing diseases [9]–[11]. Generally, these methods fall into two categories: *static FC* and *dynamic FC methods*. The *static-FC methods* characterize the FC between nodes based on the entire fMRI scan, i.e., the full time series [12], [13]. However, these methods are not equipped to characterize the dynamic properties of FCs (fluctuate over time), which are crucial for capturing the brain’s evolving states. Differently, the *dynamic-FC methods* split the whole fMRI time series into temporal segments and quantify the FC based on each segment so that time-varying FC measures can be derived. A typical pipeline for dynamic FC methods involves extracting brain network representations using GNNs for each temporal FC segment, followed by the applications of RNNs or transformers for temporal dynamics learning [4], [5], [14], [15]. Although promising, these methods still face challenges, primarily due to the unique characteristics of brain functional networks and dFC measures.

*Firstly*, **most GNN brain functional network analysis methods overlook the intrinsic brain modularity organization**, leading to suboptimal graph representation learning [9], [14], [16]–[19]. Both theoretical and empirical studies highlight that the brain functions as a modular system, comprising specialized cognitive and topological modules (also referred as functional networks in this paper). Each module or functional network comprises tightly connected brain network nodes that collaboratively perform specific functions [20], [21], as illustrated in Fig. 1(a). However, existing GNN-based methods typically treat all brain nodes uniformly, employing the same aggregation mechanism regardless of variations in node features. This limitation prevents these methods from effectively capturing modularity-specific features. *Secondly*, **most known approaches fail to capture distinctive dFC state patterns**. Evidence suggests that dFC measures often correspond to distinctive dynamic states (Fig. 1(b)), which can be identified by clustering dFC measures from temporal segments of fMRI scans [22]. Studies on brain disorders have revealed that disease-specific alterations are confined to certain dynamic states [22], underscoring that capturing dFC states can potentially improve the detection of functional brain changes associated with brain disorders.

**Fig. 1.**
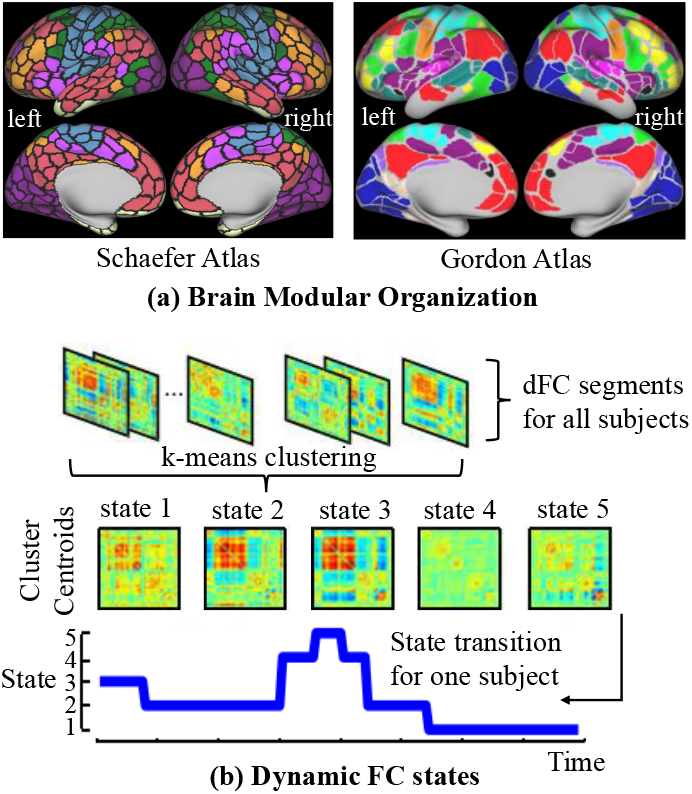
(a) The Schaefer [7] and Gordon atlases [8] with different functional networks/modules marked by different colors. Each black/grayline-bordered region represents a brain node, while regions of the same color belong to a single functional module, collectively associated with specific brain functions. (b) dFC states represent a set of recurring patterns of dFC measures, capturing the evolving FC dynamics of the brain.

To address these challenges, we propose a novel GNN-based dFC learning framework, called dFCExpert, aiming to enhance the representation learning of dynamic functional connectivity of fMRI data. The framework consists of two key components: modularity experts and state experts, specifically designed to characterize the brain modularity and the dFC state patterns. The modularity experts are built upon GNN and Mixture of Experts (MoE) for learning brain graph features of each FC segment, which characterize the brain modularity organization in the graph learning process by optimizing multiple experts at each graph layer. Each of the experts focuses on specific subsets of nodes that exhibit similar behaviors and interactions, enabling fine-grained modeling of the brain’s modular structure. On top of the graph representations learned by the modularity experts, the state experts aggregate the temporal features of dFCs into a compact set of distinctive states based on a soft prototype clustering method, where each state has similar FC patterns and reflects different activities of the dFCs related to human behaviors or brain conditions. Finally, the clustering-derived state features are used for task prediction. With the modularity and state experts, our dFCExpert introduces two novel strategies for learning-based fMRI analysis: modeling the brain functional network modularity and dynamic FC states. This pioneering approach presents a first-of-its-kind method for learning informative and explainable features of dFC patterns from fMRI data, offering significant potential for advancing imaging neuroscience research and clinical applications.

We have evaluated the performance of dFCExpert on three large-scale fMRI datasets, including the Human Connectome Project (HCP) [23], the Adolescent Brain Cognitive Development (ABCD) [24], and the Autism Brain Imaging Data Exchange (ABIDE) [25]. Across pattern recognition tasks such as classification and regression, dFCExpert consistently outperformed known methods, achieving state-of-the-art performance. To validate the framework’s design, we conducted comprehensive ablation studies to analyze the contribution of individual components of dFCExpert. Visualization results revealed that the modularity experts effectively target distinct brain modules and the state experts identify unique dynamic brain states, indicating that the method was capable of learning interpretable representations of dFC measures for prediction tasks. Moreover, we demonstrated that the brain modules and dynamic states identified by dFCExpert are biologically meaningful, bridging the gap between deep learning-based functional network modeling and traditional functional network analytic methods, highlighting the potential of dFCExpert to deliver both robust predictive performance and valuable insights into the brain’s functional organization.

## II. Related Works

### A. Dynamic Functional Connectivity Learning

To capture the time-varying FC patterns of the brain, a spatio-temporal graph convolutional network (ST-GCN) is developed to incorporate both spatial and temporal convolutions to model the non-stationary nature of dFC measures [18]. Many methods have adopted a GNN-RNN pipeline to learn graph-level features from each FC segment with the GNNs, followed by RNNs or transformers to capture temporal FC patterns [14]–[17]. Particularly, STAGIN utilizes GNNs with spatial attention and transformers with temporal attention to model the dynamics of brain networks [14]. DynDepNet introduces a dynamic graph structure learning method to capture the time-varying structures of fMRI data [17]. NeuroGraph systematically evaluates the effectiveness of different GNN designs for modeling dynamic functional networks [16]. While these methods have shown promise in modeling the dynamics of brain functional connectivity, they often overlook the brain’s intrinsic modular organization and fail to capture distinctive dFC states [14], [16]–[18]. This limitation hinders their ability to provide deeper insights into the brain’s functional organization and its evolving connectivity states.

### B. Brain Modularity Organization

The human brain functions as a modular system, composed of distinct functional and topological modules. Brain regions within the same modules are tightly connected and tend to perform similar functions [15]. For instance, salience network (SN) and default mode network (DMN) are two crucial neurocognitive modules in the brain, where the SN mainly detects internal or external stimuli and coordinates the brain’s response to those stimuli, and the DMN is responsible for self-referential thinking, mind-wandering, and introspection [20], [21]. A few existing methods have incorporated the brain modular organization into graph representation learning. BRAINNETTF introduces a novel graph readout function that leverages modular-level similarities between brain nodes to pool graph-level embeddings from clusters of functionally similar nodes [11]. MSGNN [15] develops a modularity- constrained GNN that enforces node embeddings to align with three functional network modules (i.e., central executive network, SN, DMN), by applying a modularity constraint loss after the graph layers to encourage similarity between node-level embeddings within the same module. However, since these methods incorporate the brain modularity after the graph model, all brain network nodes are still processed uniformly within each graph layer. This may be suboptimal for effectively learning representations of nodes belonging to distinct brain modules. To address this, we propose modularity experts, a combination of GNN and MoE, to mimic the brain’s modular organization, which employs multiple experts at each graph layer to guide brain network nodes with similar functions towards the same experts.

### C. Dynamic FC States

The dynamic brain states are a compact set of unique patterns derived from dFC measures, which represent connectivity patterns that repetitively occur during the fMRI acquisition [26]. Existing studies typically identify the dynamic brain states by clustering the dFC measures into different states with methods, such as k-means, and then conduct statistic analyses to investigate the relationships between these states and biological or behavior measures of interest [22]. However, this two-step strategy has significant limitation in that the features used for clustering are not explicitly optimized for the downstream statistical analyses, leading to a potential mismatch between the identified states and their informativeness for specific tasks. In contrast, our state experts overcome this limitation by adopting an end-to-end learning framework, simultaneously learning distinctive dFC states and optimizing representations of dFC graphs for specific tasks. By integrating state discovery and task optimization, our method ensures that the resulting states are both highly informative and tailored to the requirements of the analysis, enhancing their interpretability and utility in studying dFC patterns.

### D. Expert Models

The concept of Mixture of Experts (MoE) is a well-established machine learning technique that employs a divide-and-conquer strategy, where a system is composed of multiple specialized experts, each responsible for solving a specific sub- task or learning a specific substructure. Over the years, MoE has demonstrated remarkable success across various domains, including multi-modal learning [27], computer vision [28], and machine translation [29]. In the context of graph classification, for example, TopExpert applies MoE to extracted graph features, leveraging topology-specific experts for molecule property prediction [30]. More recently, a graph MoE (GMoE) model incorporates multiple experts within each graph layer to enhance the capacity of GNNs [31]. Building upon MoE, we propose modularity experts, a novel approach designed to capture the brain’s modular organization with mutilple experts. Each expert specializes in learning representations for a specific brain module, enabling our model to function in a brain-inspired manner and improving the representation learning of dFCs.

## II. Method

We begin by introducing the preliminaries of the proposed dFCExpert framework, including the concepts of GNNs and the construction of dFC graphs from fMRI data (Section III- A), followed by a detailed description of the framework, including an overview, modularity experts, state experts, and the overall loss function (Section III-B).

### A. Preliminaries

#### 1) Graph Neural Networks

To glean useful information from graph-structured data, GNNs iteratively compute node features by aggregating information of neighbor nodes and updating the node features with non-linear functions in a layer-wise manner. The propagation mechanism for a node *i* at the (*l*)-th GNN layer is formulated as:

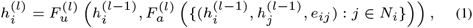

where 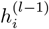 denotes the features of node *i* at the (*l* − 1) layer, *N*_*i*_ is the set of neighbors of node *i, e*_*ij*_ denotes the edge between nodes *i* and *j*, and *F*_*a*_ and *F*_*u*_ are differentiable functions for aggregating information and updating node features, respectively. Different choices of these two functions lead to various GNN architectures. For instance, Graph Isomorphism Network (GIN) [32], a variant of GNN, uses summation as *F*_*a*_ and a multi-layer perceptron (MLP) as *F*_*u*_:

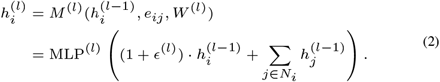

We simplify the two functions described above into a single operation *M* ^(*l*)^, where *W* ^(*l*)^ represents the trainable weight of the multi-layer perceptron (MLP), and *ϵ* is a learnable parameter initialized to zero. Given its strong ability for graph representation learning, we adopt GIN as our dFC graph feature extractor, following the approach in [14].

#### 2) Dynamic Graph Definition

To construct dFC graphs from fMRI scans, we first use a brain atlas to transform the 4D fMRI data into a time-course matrix 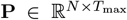. This matrix is created by averaging the fMRI signals within each brain region defined by the atlas, yielding fMRI signals of *N* brain network nodes at each time point. Next, using a sliding-window approach, the time-course matrix is divided into *T* = ⌊*T*_max_ − Γ*/S* ⌋ temporal segments, where Γ is the window length and *S* is the stride size. For each segment *t*, an FC matrix *X*_*t*_ ∈ ℝ^*N ×N*^ (*t* = 1, …, *T*) is computed by calculating Pearson correlation coefficients between the time series of all brain region pairs, yielding *T* FC matrices. Following [16], the correlation matrices can be informative node features, where the features for the *i*-th node in segment *t* correspond to the elements of the *i*-th row in *X*_*t*_. Additionally, a binary adjacent matrix *A*_*t*_ ∈ {0, 1} ^*N×N*^ is derived from *X*_*t*_ by retaining the top 30-percentile correlation values as connected and setting the remaining values as unconnected, as described in [13]. Thus, the input dFC graphs for each subject are represented as **G**_*t*_ = {*A*_*t*_, *X*_*t*_}(*t* = 1, …, *T*) (Fig. 2).

**Fig. 2.**
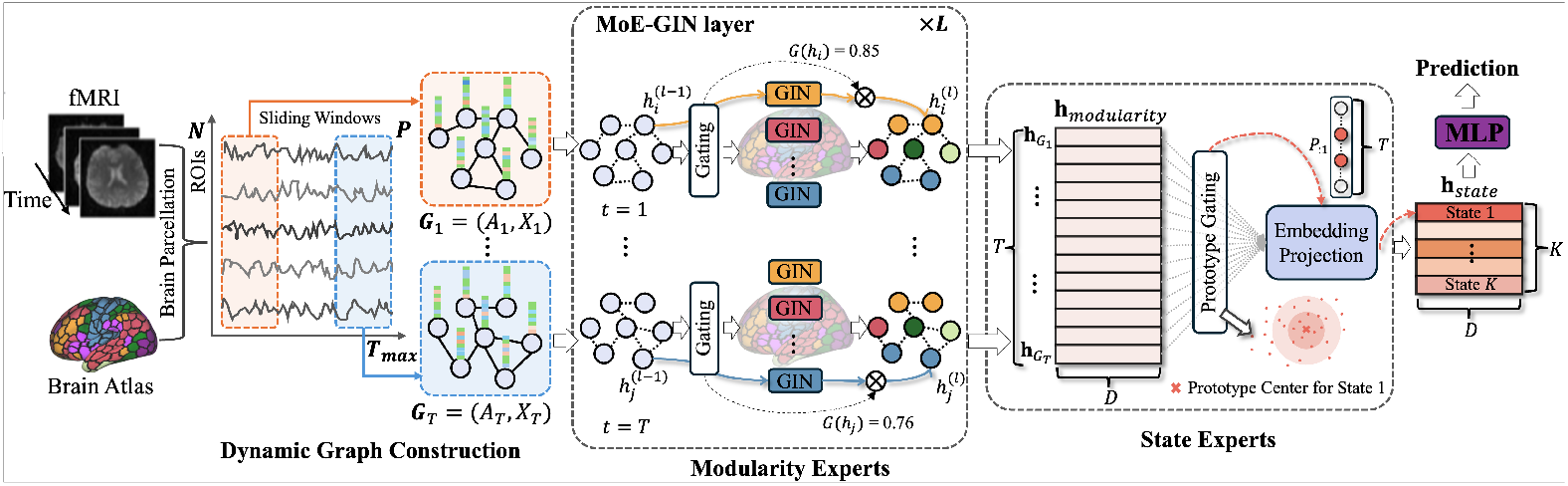
The dFCExpert framework consists of modularity experts and state experts. The modularity experts include ***L*** layers of MoE-GIN, which route node features ***h***_***i***_ to a specific GIN expert with a gating score ***G*(*h***_***i***_) for learning graph-level features 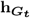 for each temporal segment ***t***, yielding 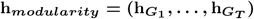 across all ***T*** segments. The state experts aggregate the learned temporal graph features into dynamic states using prototype gating and embedding projection. Finally, the state features **h**_***state***_ are processed through an MLP layer for task prediction.

### B. dFCExpert

#### 1) Overview

As illustrated in Fig. 2, dFCExpert consists of modularity experts and state experts. Taking the dFC graphs as input, the modularity experts leverage a combination of GIN and MoE to learn brain graph features for each FC segment, aiming to capture the brain modularity mechanism effectively. Building upon the outputs of the modularity experts, the state experts adaptively group the temporal graph features into distinctive states using a soft prototype clustering method. This approach allows the model to learn expressive state features by assigning soft clusters to the temporal graph features. Finally, the learned state features are passed through an MLP layer to predict a specific task, such as predicting sex in a classification setting or predicting an intelligence measure in a regression setting. Formally, the objective of dFCExpert is to train a neural network *f* : (**G**_1_, …, **G**_*T*_) → **h**_*state*_, where **G**_*t*_ = {*A*_*t*_, *X*_*t*_}(*t* = 1, …, *T*) represents the sequence of constructed dFC graphs with *T* segments, and **h**_*state*_ ∈ ℝ^*K×D*^ is the output features from the state experts. We formulate *f* = *s*∘*m* as a composition of modularity experts *m* and state experts *s*, where *m* learns dFC graph representations **h**_*modularity*_ = 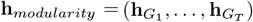 for each of *T* temporal segments, and *s* aggregates these learned temporal graph features into the final state features **h**_*state*_:

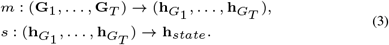

#### 2) Modularity Experts

The modularity experts are designed to emulate the human brain’s modular organization, enabling effective learning of brain graph representations. The modularity experts are implemented using a combination of GIN and MoE, referred to as MoE-GIN, consisting of *L* layers of MoE-GIN. Specifically, each MoE-GIN layer includes one gating function and multiple GIN experts, and all *T* segments share the same MoE-GIN layer, as illustrated in Fig. 2. The gating function determines which experts are most suitable for processing a given node, while the multiple experts are independent GIN propagation functions (Eq. (2)), each with its own trainable parameters. This design of MoE-GIN enables nodes that are tightly connected or behaviorally similar to be routed to the same expert. Consequently, each GIN expert specializes in learning representations for a specific brain module to effectively capture distinct brain activities. By organizing the brain nodes into specialized groups, the modularity experts mimic the way the brain operates, where tightly connected regions interact to perform specific functions or activities. This design allows the model to reflect the brain’s modularity, improving its ability to learn meaningful and interpretable representations of brain networks. Formally, the operation of a single MoE-GIN layer is described as:

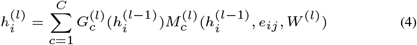

where *C* is the number of experts, and *M* denotes the node propagation function of GIN as defined in Eq. (2). *G* is a gating function to generate assignment scores based on the input of node features 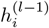, and 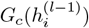 denotes the score for the *c*-th expert. We implement a simple gating function by multiplying the input with a trainable weight matrix *W*_*g*_, followed by the application of the softmax function [33]: *G*(*h*_*i*_) = softmax(*h*_*i*_*W*_*g*_). Given that each node typically belongs to a single brain module, we adopt a simplified top-1 gating scheme as in [34], where each node is routed to only one expert, corresponding to the expert with the highest gating score. This simplification reduces computational overhead of routing while maintaining model quality and interpretability. Additionally, each of the *T* segments will go through the *L* layers of MoE-GIN, we omit segment notation *t* in Eq. (4) for brevity, whenever it is not of contextual importance.

The graph-level representation 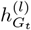 for segment *t* at layer *l* is computed by averaging the updated node features 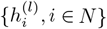, and the final graph representation 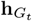 is then obtained by concatenating the graph representations from all *L* layers [35] followed by an MLP layer for dimension reduction: 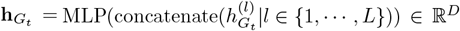. Finally, we obtain 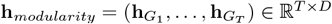.

##### Auxiliary Loss Functions for Training MoE-GIN

If the model is trained solely using the task-specific prediction loss, the gating network may converge to trivial solutions where only a few experts are consistently selected [36]. This imbalance in expert selection becomes self-reinforcing, as the favored experts dominate the learning process, further increasing their frequency of selection. Another trivial situation is that the gating network may produce similar assignment scores across all experts for a given node, resulting in a lack of specialization in experts. In our scenario, we expect the gating network to be capable of effectively distinguishing nodes of different brain functional modules and enable sparse gating so that the selected expert receives a significantly higher score than the others. To achieve this, we introduce two auxiliary loss functions to promote balanced loading and enforce sparse gating, respectively. Given a batch *B*, each containing *T* graphs, the loading balance loss is computed as:

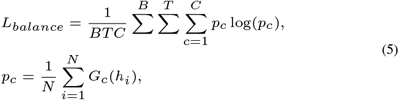

where *N* is the number of nodes, and *p*_*c*_ represents the fraction of nodes routed to expert *c*. By minimizing the negative entropy of the *p*_*c*_ distribution, the *L*_*balance*_ encourages a uniform distribution of nodes across all experts. To promote sparse gating in *G*(*h*_*i*_), we introduce a sparse gating loss:

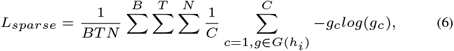

where *g*_*c*_ represents the gating score for assigning a given node to expert *c*. Minimizing the entropy of the gating scores across all experts encourages sparsity, ensuring each node is strongly associated with a specific expert. We implement these two loss functions similar to [37], as they are both fundamentally deep clustering-based and exert opposing influences during training.

#### 3) State Experts

The state experts follow the traditional dFC analysis in imaging neuroscience to adaptively group the temporal graph features **h**_*modularity*_ into a small number of states, each reflecting distinct dynamic states of dFC measures associated with different brain conditions. From another perspective, representing dFC measures with a smaller set of states simplifies the model, reduces data complexity, and makes it easier to interpret the FC dynamics. Specifically, we design the state experts using a soft prototype clustering method, where temporal FC graph features are softly assigned to clusters in an unsupervised manner. As illustrated in Fig. 2, distinctive dFC states are characterized through prototype gating and embedding projection. The prototype gating introduces *K* trainable prototype centroids *U* = [*u*_1_, …, *u*_*K*_], each with *D* dimensions (*U* ∈ ℝ^*K×D*^), to adaptively learn the feature centers of each state. Given the learned graph features 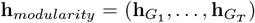, the prototype gating calculates the assignment score *P*_*tk*_ for assigning segment *t* to state *k* using a Softmax projection:

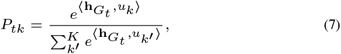

where ⟨*·,·*⟩ denotes the inner product. The embedding projection aggregates the temporal graph features **h**_*modularity*_ ∈ ℝ^*T ×D*^ into state features **h**_*state*_ under the guidance of the obtained soft assignment score *P* ∈ ℝ^*T ×K*^. Specifically, to compute features for a single state (e.g., State 1 as in Fig. 2), the embedding projection integrates features of the temporal segments assigned to that state using soft gating, performed as 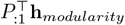. This process ensures that the state features are derived by aggregating the temporal FC features with high probabilities assigned to a specific state, thus enabling the model to effectively capture the dynamic nature of FC patterns and associate them with interpretable states.

Since the clustering lacks explicit guidance labels, we optimize the temporal FC features and cluster centroids in a self-supervised manner by leveraging orthonormal initialization of *U* [11] and optimizing the cluster distribution for cohesive clustering. Specifically, we define a target distribution *Q*_*tk*_ for each 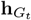 based on the current cluster assignment distribution *P*_*tk*_, which therefore enhances high-confident assignments [38] through a squaring operation, defined as:

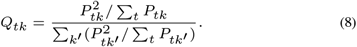

This target distribution strengthens the cluster cohesion, refining temporal FC representations to become more discriminative with respect to their state patterns. Finally, we minimize a KL divergence loss to align *P* and *Q*, boosting the cohesion of the clusters:

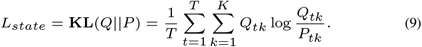

#### 4) Overall Loss Function

The final loss combines a task-specific loss (*L*_*task*_), two auxiliary losses for the modularity experts (*L*_*balance*_ and *L*_*sparse*_), and a loss for state experts (*L*_*state*_), yielding the overall optimization objective. Two scaling factors, *α* and *β*, are used to balance the contributions of these components:

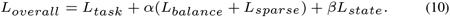

After experimenting with various scaling ratios, we set *α* = 1 and *β* = 10 for optimal performance. Note that the auxiliary losses for modularity experts are computed as the summation across all MoE-GIN layers.

## IV. Experiments

### A. Experimental Settings

#### 1) Datasets

We conducted experiments on three fMRI datasets. (a) *Human Connectome Project (HCP)*: We used data from the HCP S1200 release [23], which includes preprocessed and ICA de-noised [39] resting-state fMRI scans of 1071 individuals with 578 females and 493 males. Following [14], we selected data from the first run and excluded scans with fewer than 1200 time points. All the resting-state fMRIs used in this study contain exactly 1200 time points. To build functional networks, we parcellated the brain into 400 regions using the Schaefer atlas [7], which is organized into seven intrinsic connectivity networks. (b) *Adolescent Brain Cognitive Development Study (ABCD)*: The study is a multi-site investigation of brain development and its relationship with behavioral outcomes in children aged 9-10 years [24]. In our study, we used preprocessed baseline resting-state fMRI data from the ABCD BIDS Community Collection (ABCC) [40]. As in [41], participants with incomplete data, excessive head motion, or fewer than 600 remaining time points after motion censoring were excluded. After quality control, our final dataset included resting-state fMRI scans of 6,195 children (3,080 females and 3,115 males), with the number of time points ranging from 626 to 3,516. The Gordon atlas [8] with 352 regions (*N* = 352) was used for brain parcellation, as defined by the ABCC. (c) *Autism Brain Imaging Data Exchange (ABIDE)*: The ABIDE initiative aggregates functional brain imaging data from multiple sites to support research on Autism Spectrum Disorder (ASD) [25]. We used data from [42], comprising fMRI scans from 1025 subjects (537 typical controls and 488 individuals with ASD), with time points ranging from 120 to 300. Similar to HCP, the Scheafer atlas with 400 regions was used for brain parcellation.

All datasets are publicly available, with all identification information anonymized. The image acquisition parameters for the HCP, ABCD and ABIDE datasets can be found in [39], [24], and [43], respectively. We created 5 stratified splits for all datasets with a train-validation-test ratio of 7:1:2. Average results and standard deviations across the 5 splits were reported.

#### 2) Targets and Metrics

We selected three evaluation tasks: sex and ASD classification, and cognitive intelligence prediction (“fluid intelligence” for HCP and “general cognition” for ABCD). The sex/ASD classification task is a binary classification problem, and the cognitive intelligence prediction is a regression problem. These tasks were chosen to explore brain-biology and brain-cognition associations. For the sex/ASD classification task, we used accuracy (ACC) and AUC (Area Under ROC Curve) as evaluation metrics. For the regression task, the regression targets were z-normalized and the performance was evaluated using Mean Squared Error (MSE) and correlation coefficient (CORR) between the measured and predicted values.

#### 3) Implementation Details

Our method was implemented using PyTorch [44] and trained on an NVIDIA A100 GPU with 80 GB memory. We set the number of MoE-GIN layers to *L* = 3 and the embedding dimension to *D* = 256. For Dfc graph construction, we used a window length of Γ = 50 and a window stride of *S* = 3, following common protocols for sliding-window-based dFC analyses [26], [45]. Consistent with [14], we randomly selected 600 time points at each training step for the dFC graph computation. This procedure reduced computational and memory overhead while augmenting the training data. For testing, we utilized the full time-course matrix 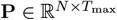 to construct dFC graphs and evaluate the model. The sex/ASD classification and cognitive intelligence regression tasks were trained using cross-entropy and MSE losses, respectively. To ensure a fair comparison, we trained both dFCExpert and known methods under comparison in an end-to-end supervised fashion using a similar configuration:

- *Optimizer*: Adam.
- *Learning rate*: A learning rate of 5*e*^−4^ for training the classification task, and a learning rate of 1*e*^−3^ for training the regression task.
- *Mini-batch size*: mini-batch size of 8 for the HCP/ABIDE dataset and mini-batch size of 16 for the ABCD dataset.
- *Epochs*: 50 epochs for the HCP/ABIDE dataset, and 30 epochs for the ABCD dataset.
- *Number of experts*: *C* = 7 for HCP/ABIDE dataset and *C* = 5 for ABCD dataset.
- *Number of states*: *K* = 7 for HCP/ABIDE dataset and *K* = 5 for ABCD dataset.

Based on our experiments, *C* can be set to 5 or 7, and *K* can range from 5 to 7. These values for *C* and *K* fall within a reasonable and effective range for our model’s performance.

### B. Sex Classification and Cognitive Intelligence Regression Results

#### 1) Comparison with Existing Methods

We evaluated the performance of dFCExpert against alternative *static-* and *dynamic-FC methods* on all the sex/ASD classification and cognitive intelligence regression tasks. The results of these alternative methods are summarized in the top two blocks of Table I. For the sex classification task, dFCExpert outperformed all the alternative methods under comparison on both the HCP and ABCD datasets. On the regression task, our method demonstrated significant performance improvements, particularly on the HCP dataset, with gains of nearly 7 points. For ASD classification on a disease dataset, dFCExpert also achieved superior performance. These quantitative results highlight the importance and effectiveness of incorporating brain modularity and learning state patterns to model brain dynamics in fMRI data, underscoring the advantages of our approach.

**TABLE I.** Performance comparison with alternatives and baselines. (“M”= Modularity, “S” = State)

| Method | HCP |  |  |  | ABCD |  |  |  | ABIDE |  | FC Type |
| --- | --- | --- | --- | --- | --- | --- | --- | --- | --- | --- | --- |
|  | Sex |  | Intelligence |  | Sex |  | Cognition |  | ASD |  |  |
|  | ACC(%) | AUC(%) | MSE(↓) | CORR(↑) | ACC(%) | AUC(%) | MSE(↓) | CORR(↑) | ACC(%) | AUC(%) |  |
| BrainNetCNN [10] | 84.56±1.87 | 91.97±1.33 | 1.130±0.039 | 0.200±0.071 | 85.38±0.07 | 92.67±0.33 | 0.474±0.031 | 0.442±0.011 | 64.29±1.73 | 72.66±1.93 | Static |
| BRAINNETTF [11] | 85.42±2.36 | 92.51±2.01 | 1.015±0.052 | 0.205±0.074 | 84.55±0.64 | 91.38±0.21 | 0.462±0.022 | 0.465±0.024 | 65.57±2.06 | 70.13±1.61 | Static |
| NeuroGraph [16] | 84.38±2.60 | 92.03±2.39 | 1.321±0.043 | 0.198±0.065 | 85.71±0.60 | 93.75±0.61 | 0.512±0.013 | 0.423±0.015 | 64.98±3.19 | 70.94±2.53 | Static |
| STAGIN [14] | 87.74±1.75 | 92.09±0.93 | 1.002±0.042 | 0.215±0.078 | 86.93±0.45 | 91.57±0.57 | 0.452±0.012 | 0.477±0.008 | 60.86±4.30 | 67.24±4.28 | Dynamic |
| MSGNN [15] | 86.25±2.03 | 92.37±1.42 | 1.073±0.057 | 0.206±0.070 | 87.13±0.63 | 92.67±0.42 | 0.453±0.024 | 0.483±0.017 | 65.36±3.16 | 70.46±3.42 | Dynamic |
| NeuroGraph [16] | 84.66±3.30 | 93.14±2.23 | 1.017±0.032 | 0.207±0.069 | 87.37±1.08 | 94.64±0.69 | 0.441±0.031 | 0.482±0.005 | 65.18±3.04 | 70.28±3.36 | Dynamic |
| Baseline | 88.11±4.26 | 96.68±1.71 | 0.978±0.079 | 0.238±0.086 | 87.25±0.70 | 94.86±0.70 | 0.442±0.021 | 0.484±0.013 | 65.85±3.12 | 72.46±2.96 | Dynamic |
| Baseline + M | 89.61±4.22 | 96.91±1.68 | 0.946±0.041 | 0.272±0.059 | 88.09±0.76 | 95.16±0.54 | 0.425±0.026 | 0.499±0.018 | 67.29±3.05 | **74.87±3.30** | Dynamic |
| Baseline + S | 89.98±2.83 | 97.23±1.32 | 0.967±0.074 | 0.248±0.083 | 88.56±1.25 | 95.81±0.53 | 0.419±0.014 | 0.496±0.013 | 66.17±3.05 | 74.31±2.82 | Dynamic |
| dFCExpert | **91.03±1.71** | **97.40±1.30** | **0.932±0.036** | **0.296±0.049** | **89.28±0.83** | **95.99±0.52** | **0.412±0.012** | **0.513±0.009** | **68.93±2.89** | 74.62±3.02 | Dynamic |

#### 2) Ablation Studies

We conducted ablation experiments to evaluate the contributions of individual components of dFC- Expert, specifically the modularity experts and state experts. For fair comparison, the baseline model was a 3-layer GIN followed by an MLP layer. The GIN was used to learn graph-level features, and the MLP made predictions for each segment *t*, which were then averaged to generate the final result. The baseline model was chosen for the following reasons: 1) Generating predictions for each temporal segment provides stronger supervision, potentially improving prediction compared to the alternatives that use RNNs or transformers on top of the GIN layer (e.g., STAGIN [14] and NeuroGraph [16], as shown in Table I); 2) Since the state experts aggregate the temporal graph features into *K* state features, the use of RNNs or transformers for final predictions is unnecessary. Based on this baseline, we introduced modularity experts (“Baseline + M”) by replacing GIN with MoE-GIN, and state experts (“Baseline + S”) by aggregating temporal features into several connectivity states before applying the MLP layer, respectively. The third block of Table I summarizes the ablation study results, from which several key observations can be made: 1) Compared with the baseline model, the modularity experts achieved significantly improved performance, particularly for the regression task on the HCP dataset, highlighting the effectiveness of characterizing brain modularity for learning more informative brain graph features; 2) By grouping the temporal FC measures into several states, the state experts achieved better performance across both tasks and both datasets, underscoring the benefit of using this new approach to explore the dynamics of FC measures; 3) Incorporating both the modularity and state experts, dFCExpert outperformed the baseline model, validating the overall effectiveness of our method. These results demonstrate that both components play a crucial role in improving the model’s ability to capture dynamic brain connectivity and its association with cognitive and biological outcomes.

### C. Modularity Experts Analysis

We evaluated the effectiveness of our modularity experts from the following perspectives: 1) the impact of the number of experts in the MoE-GIN layer; 2) the ability of modularity experts to learn specific brain modules; 3) the performance of modularity experts when applied to *static-FC* measures; and 4) the effect of different scaling ratios for the auxiliary losses.

#### 1) Different Number of Experts

To evaluate the effect of modularity experts on model performance, we investigated the impact of the number of experts (*C*) in the MoE-GIN layer. Specifically, we tested *C* values of 3, 5, 7, 9, 17, where *C* = 7 and *C* = 17 are widely used settings in large-scale functional network studies [46]. The results are summarized in Table II and reveal the following insights: 1) On the HCP dataset, the model’s performance improved as the number of experts increased from 3 to 7 but began to decline when the number of experts increased further from 7 to 17, suggesting that a smaller number of experts is not only optimal for performance but also beneficial for reducing computational cost and model complexity. 2) On the ABCD dataset, the modularity experts outperformed the baseline across all settings. While the number of experts had a smaller effect on the classification task, the performance on the regression task improved significantly as *C* increased, peaking at *C* = 9. 3) On the ABIDE dataset, we observed similar trends as in the HCP dataset, with *C* = 7 yielding the best performance for ASD identification. These results suggest that *C* = 5 or *C* = 7 are reasonable choices, consistent with the typical number of functional brain modules. We selected *C* = 7 for the HCP and ABIDE datasets, and *C* = 5 for the ABCD dataset. The variation in optimal number of experts likely reflects differences in brain parcellation strategies and the intrinsic characteristics of each dataset, as HCP (young adults), ABCD (young kids) and ABIDE (disease) represent different age groups, developmental stages, and health conditions.

**TABLE II.** Impact of the number of experts in the modularity experts on performance.

| #Experts | HCP |  |  |  | ABCD |  |  |  | ABIDE |  |
| --- | --- | --- | --- | --- | --- | --- | --- | --- | --- | --- |
|  | Sex |  | Intelligence |  | Sex |  | Cognition |  | ASD |  |
|  | ACC(%) | AUC(%) | MSE(↓) | CORR(↑) | ACC(%) | AUC(%) | MSE(↓) | CORR(↑) | ACC(%) | AUC(%) |
| 1 (baseline) | 88.11±4.25 | 96.68±1.71 | 0.978±0.079 | 0.238±0.086 | 87.25±0.70 | 94.86±0.70 | 0.442±0.021 | 0.484±0.013 | 65.85±3.12 | 72.46±2.96 |
| 3 | 88.68±2.57 | 96.46±1.58 | 0.961±0.067 | 0.266±0.061 | 87.57±0.94 | 95.16±0.70 | 0.435±0.019 | 0.481±0.005 | 66.02±3.11 | 73.65±3.70 |
| 5 | 89.25±3.46 | 96.41±1.27 | 0.953±0.054 | **0.279±0.042** | **88.09±0.76** | **95.16±0.54** | **0.425±0.026** | **0.499±0.018** | 66.90±3.98 | 73.18±2.94 |
| 7 | **89.61±4.22** | **96.91±1.68** | **0.946±0.041** | 0.272±0.059 | 87.80±1.06 | 94.85±0.65 | 0.429±0.023 | 0.481±0.016 | **67.29±3.05** | **74.87±3.30** |
| 9 | 88.73±3.48 | 95.26±2.10 | 0.963±0.036 | 0.224±0.051 | 87.65±0.87 | 94.98±0.69 | **0.425±0.020** | 0.479±0.010 | 66.61±3.52 | 73.41±3.61 |
| 17 | 87.83±3.83 | 95.87±1.47 | 0.976±0.032 | 0.209±0.069 | 87.19±1.30 | 94.84±0.53 | 0.431±0.024 | 0.477±0.020 | 65.58±3.56 | 72.25±3.70 |

#### 2) Visualization of the Learned Brain Modules

To further evaluate the effectiveness of the modularity experts, we visualized the node assignment results produced by the modularity experts with *C* = 7 on HCP dataset and *C* = 5 on ABCD dataset, and compared them with the Schaefer Atlas (7 networks) and Gordon Atlas (12 networks) (Fig. 3 (a)), respectively. Specifically, given the gating score of last MoE-GIN layer of modularity experts, we first average the scores for all *T* segments and then average through all subjects (Fig. 3 (b)) or randomly choose one male and female subjects (Fig. 3 (c)(d)). Finally, we perform *argmax* operation on the resulting scores, and we can obtain 7 and 5 learned modules from modularity experts for HCP and ABCD datasets, respectively, where each module consists of nodes assigned to a specific expert. For the Gordon atlas, we simply visualize the 5 learned modules, as shown the second row of Fig. 3. For the Schaefer atlas, we perform a color match. Since the correspondence between these expert-learned modules and the 7 atlas networks was unknown, we matched them based on the maximum overlap of their respective nodes using the Hungarian method [47], and the matched results are visualized in the first row of Fig. 3. We can observe that the visualization demonstrates a strong overlap between the atlas networks and the modules learned by the modularity experts, with a Dice coefficient of 0.71 and *p <* 0.001, as determined by a spin-based spatial permutation test [48]). This result indicates that our modularity experts effectively grouped tightly connected nodes into coherent modules, enabling each expert to specialize in capturing the structure of a specific brain functional module. Moreover, because the modularity experts were optimized for specific tasks, they could potentially learn node-expert (or node-network) assignments that go beyond the prior knowledge encoded in the Atlases. From Fig. 3 (c)(d), we observed individual differences in resulting learned modules across subjects, indicating the ability of our modularity experts to learn unique FC patterns of each individual.

**Fig. 3.**
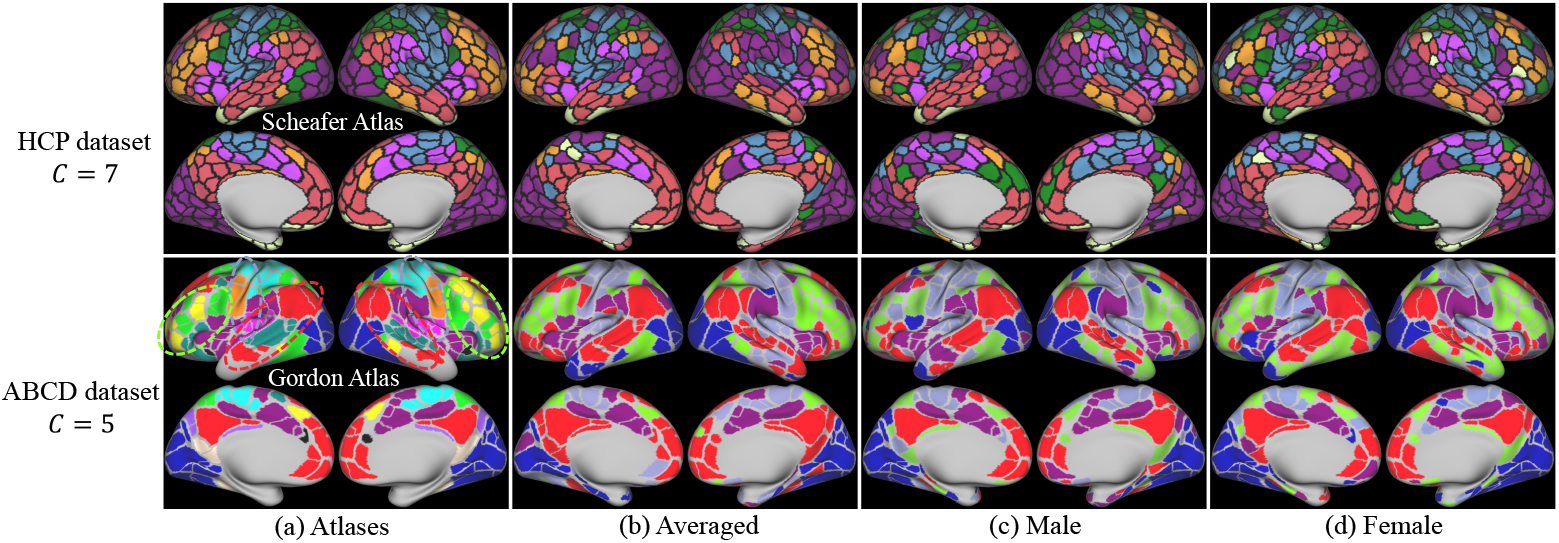
Visualization of the learned brain modules (node assignment results) averaged across all subjects (b), and for two randomly selected individuals (c) and (d) from the test datasets of HCP (first row) and ABCD (second row). These are compared to the functional networks defined by the Schaefer and Gordon atlases (a). In the first row, the seven functional networks from the Schaefer atlas are color-matched with our learned modules (***C* = 7**). In the second row, since the number of learned modules (***C* = 5**) does not match the 12 networks in the gordon atlas, we highlight key networks using consistent colors: blue for the vision network, red for the default network, and purple for the cingulo opercular network in (b)(c)(d). Other combined modules are represented using lilac and bright green.

#### 3) Modularity Experts in Static-FC Method

To investigate the effectiveness of modularity experts in static FC analysis, we integrated the MoE-GIN layer into Neurograph framework [16] that employs a GNN architecture with residual connections and combines hidden representations from message passing at each layer by concatenation. These combined representations are then processed by an MLP layer for predictions. In Table III, incorporating the modularity experts into a *static-FC* method enhanced brain graph representations and improved performance, particularly on the HCP dataset. This highlights the versatility of modularity experts in various FC analyses.

**TABLE III.** Performance of modularity experts (MoE-GIN) APPLIED TO **STATIC-FC** MEASURES.

| Method | GNN Type | HCP |  |  |  | ABCD |  |  |  |
| --- | --- | --- | --- | --- | --- | --- | --- | --- | --- |
|  |  | Sex |  | Intelligence |  | Sex |  | Cognition |  |
|  |  | ACC(%) | AUC(%) | MSE(↓) | CORR(↑) | ACC(%) | AUC(%) | MSE(↓) | CORR(↑) |
| NeuroGraph [16] | GIN | 84.38±2.60 | 92.03±2.39 | 1.321±0.043 | 0.198±0.065 | 85.71±0.60 | 93.75±0.61 | 0.512±0.013 | 0.423±0.015 |
| NeuroGraph [16] | MoE-GIN | **87.30±1.33** | **95.56±1.68** | **0.989±0.035** | **0.243±0.052** | **87.08±0.74** | **93.12±0.52** | **0.486±0.021** | **0.477±0.017** |

#### 4) Scaling Ratio α Analysis for Auxiliary Losses

We analyzed the impact of different scaling ratios *α* for the two auxiliary losses in the modularity experts on the HCP dataset, experimenting with *α* ∈ {0.1, 1}. The results are summarized in Table IV. When *α* = 0.1, performance improved on both tasks, highlighting the importance of the auxiliary losses for balance expert loading and sparse gating. With *α* = 1, classification performance further improved, although the enhancement on the regression task was less significant. For consistency, we used *α* = 1 in all subsequent experiments. To further validate the effectiveness of the proposed loading balance loss *L*_*balance*_ and sparse gating loss *L*_*sparse*_, we conducted experiments by removing either *L*_*balance*_ or *L*_*sparse*_. The results showed that the model’s performance decreased, underscoring their role in enabling balanced selection and specialization of experts.

**TABLE IV.** Performance of modularity experts with different scaling ratios ***α*** (***αb*** AND ***αs*** REPRESENT THE SCALING RATIOS FOR ***L***_***balance***_ AND ***L***_***sparse***_, RESPECTIVELY).

| $\alpha$ | $\alpha_b, \alpha_s$ | Sex | | Intelligence | |
| --- | --- | --- | --- | --- | --- |
|  |  | ACC(%) | AUC(%) | MSE(↓) | CORR(↑) |
| 0 | $\alpha_b = 0, \alpha_s = 0$ | 88.56±3.45 | 94.67±1.68 | 0.962±0.022 | 0.256±0.048 |
| 0.1 | $\alpha_b = 0.1, \alpha_s = 0.1$ | 89.16±4.12 | 95.52±2.01 | 0.948±0.032 | 0.270±0.036 |
| 1 | $\alpha_b = 1, \alpha_s = 1$ | **89.61±4.22** | **96.91±1.68** | **0.946±0.041** | **0.272±0.059** |
| - | $\alpha_b = 1, \alpha_s = 0$ | 87.83±4.05 | 94.84±2.19 | 0.965±0.028 | 0.252±0.050 |
| - | $\alpha_b = 0, \alpha_s = 1$ | 88.65±3.52 | 95.13±1.53 | 0.956±0.023 | 0.261±0.043 |

### D. State Experts Analysis

To assess the impact of prototype learning for identifying dFC states, we conducted experiments on HCP dataset to explore the performance of different numbers of states, visualize how these learned states provide insights for interpretability, and analyze the effect of the scaling factor *β* for *L*_*state*_.

#### 1) Different Numbers of States

We examined the effect of varying the number of states *K* within the range of 3, 5, 7, 9, 10, and 20. Fig. 4 (d) shows how the classification and prediction performance changed with the number of states. Consistent with observations from the modularity analysis, performance initially improved as *K* increased but began to degrade beyond a certain number of states. The best performance was achieved with *K* = 7 for the classification task and *K* = 5 for the regression task. This also aligns with the findings in existing dFC studies, which suggest that the typical number of dynamic states ranges from 5 to 7 [22].

**Fig. 4.**
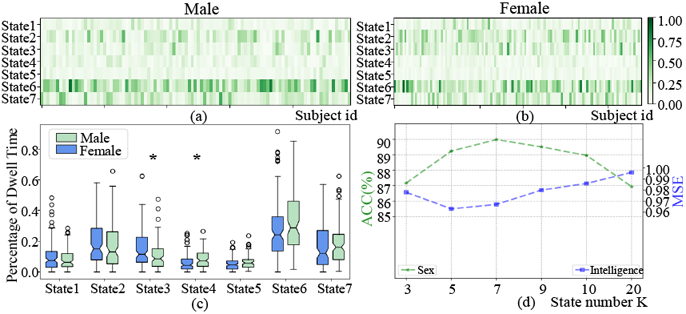
**a)** and **b)** Probabilities of temporal segments of individual males and females assigned to specific states, with the *x* -axis representing individuals in the testing dataset; **c)** Boxplots summarizing **a)** and **b)**, where each box corresponds to a row in **a)** or **b)**, and ***⋆*** indicates statistically significant difference; **d)** Model performance with different numbers of the dFC states.

#### 2) Visualization of Learned dFC States

We visualized the state assignment results in Fig. 4 and the averaged FCs of temporal segments in Fig. 5 to demonstrate whether the learned states provide interpretable evidence for distinguishing males from females. Specifically, we averaged the soft assignment results *P* across the temporal dimension *T*, such that each resulting value represented the proportion of a subject’s temporal segments assigned to a specific state (e.g., the *k*-th state), where each state captures a representative, summary patterns of the dFC measures. These proportions were computed for all subjects in the HCP test dataset and visualized separately for males and females. The results shown in Fig. 4 (a) and (b), along with the boxplot in Fig. 4 (c) reveal distinct patterns. From Fig. 4 (a) and (b), we observe that female dFC graphs are more frequently assigned to States 2 and 3 compared to male dFC graphs. Although both groups show frequent assignments to State 6, males exhibit a higher possibility. In contast, States 1, 5, and 7 show no significant differences in assignment between males and females. These findings are further supported by the boxplot in Fig. 4 (c), where notable differences are observed in States 3, 4 and 6, particularly States 3 and 4. Wilcoxon rank-sum tests indicate statistically significant differences between males and females in these states, with *p*-values less than 0.05. This indicates that sex differences are most pronounced in States 3 and 4, with State 3 being more characteristic of females and State 4 of males. Overall, the different state assignment patterns between females and males suggest that the learned dFC states capture meaningful and interpretable sex-related differences. These results also highlight the potential clinical relevance of clustering-derived dFC states, demonstrating that male and female groups are differentially associated with specific dFC states.

**Fig. 5.**
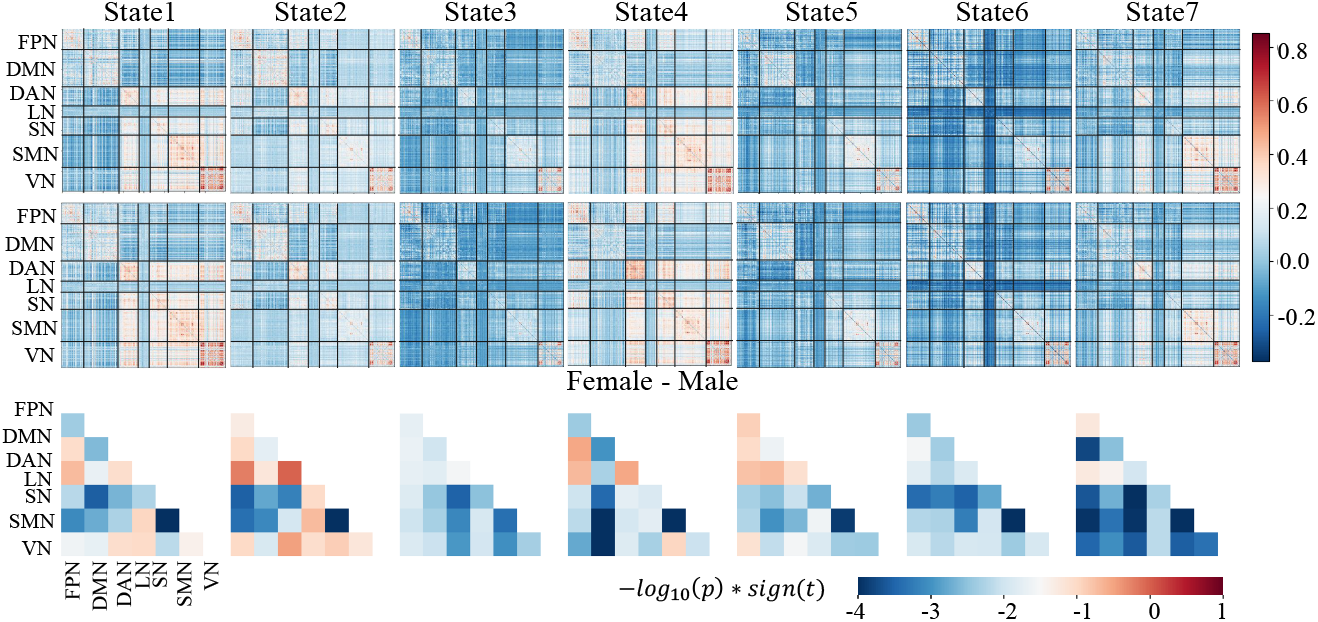
Visualization of the averaged FC measures across temporal segments for males and females assigned to each specific state. The first row shows results for the female group, the second row displays results for the male group, and the third row highlights sex differences in FC. These differences were assessed using a two-sample ***t***-test comparing the averaged FCs between female and male groups for each state. Brain network nodes were grouped into seven functional networks. The visualized differences are represented as **−*log***_**10**_**(*p*) *× sign*(*t*)**, where ***p*** and ***t*** are the ***p***-value and ***t***-statistic from the two-sample ***t***-test. For reference, ***log***_**10**_**(0.05) *≈* −1.30**. The y-axis corresponds to the seven networks: Frontoparietal Network (FPN), Default Mode Network (DMN), Dorsal Attention Network (DAN), Limbic Network (LN), Salience Network (SN), Somatosensory Motor Network (SMN), and Visual Network (VN).

To examine how brain regions are specifically connected within the learned states and how they differ by sex, we visualized the averaged FCs of temporal segments assigned to each state for males and females. As shown in the first two rows of Fig. 5, each state exhibits distinct connectivity patterns, adaptively summarized from the temporal FC measures. In these visualizations, red indicates strong positive connectivity (correlation), while blue indicates strong negative connectivity. For example, in States 1 and 4, the DAN shows strong positive connectivity with the SN, SMN and VN. In contrast, the LN shows strong negative connectivity with all other networks in State 6. These distinct patterns across states validate the effectiveness of our state experts in aggregating similar FC patterns and capturing meaningful brain dynamics. From the female-male difference, we find that sex-related differences in FC are particularly evident in the SN and SMN, where significant differences between male and female groups are observed across nearly all states. These findings are consistent with prior studies on sex differences in functional networks [49]. Specifically, in States 1, 2, and 3, significant sex differences are observed in the connectivity between SN and DMN, SN and FPN, and SN and DAN, respectively. In State 4, significant connectivity differences appear in DMN, SMN and VN. In State 7, almost all networks exhibit significant sex-related differences, especially in the connectivity between DAN and FPN, SMN and FPN, SN and DAN, and SMN and DAN. Consistent with the state assignment results in Fig. 4, the FC patterns in States 3 and 4 also show pronounced sex differences, reinforcing the interpretability of these states. These findings demonstrate that the learned dFC states capture explainable and biologically meaningful sex differences in dFC. Moreover, they highlight the ability of our state experts to identify distinctive dFC states that align with findings from conventional dFC studies [22], [26], despite employing a different learning strategy.

Overall, the observed sex differences in both state assignment and FC strength underscore the interpretability and clinical relevance of our state experts, emphasizing their capacity to capture meaningful variations in dFC.

#### 3) Scaling Factor β Analysis for L_state_

We investigated the impact of the scaling factor *β* for the clustering loss *L*_*state*_ on the HCP dataset. As shown in Table V, *β* = 1 resulted in improved performance compared to *β* = 0, demonstrating the critical role of the clustering loss *L*_*state*_ in learning meaningful state patterns. Since *L*_*state*_ is relatively small in magnitude, we also tested a larger scaling factor, *β* = 10, to amplify its influence. The resulting performance showed further improvement, indicating enhanced state assignment.

**TABLE V.** Results for state experts with different scaling ratios ***β***.

| $\beta$ | Sex | | Intelligence | |
| --- | --- | --- | --- | --- |
| | ACC(%) | AUC(%) | MSE(↓) | CORR(↑) |
| 0 | 86.75±2.43 | 93.52±1.12 | 1.023±0.065 | 0.210±0.072 |
| 1 | 88.23±2.12 | 96.28±1.85 | 0.974±0.044 | 0.268±0.063 |
| 10 | **89.98±2.83** | **97.23±1.32** | **0.967±0.074** | **0.248±0.083** |

## V. Conclusions and Discussions

We developed dFCExpert, a novel method for learning effective representations of dFC measures, which consists of modularity experts and state experts, designed to reflect the brain’s modular organization and dFC states that have been extensively studied in fMRI research on functional neuroanatomy, brain development and aging, and brain disorders, but underexplored in the machine learning community for functional brain networks. As demonstrated by extensive experiments on three large-scale fMRI datasets, the modularity experts automatically routed network nodes with similar brain functions to the same expert, enabling specialization of experts and promoting effective representation learning of dFC measures. Meanwhile, the state experts grouped the learned temporal graph features into distinctive states, each characterized by similar dFC patterns. This facilitated the effective modeling of temporal dynamics in brain functional networks, revealing insights into different brain states and biological characteristics. These results highlight not only the superior performance of dFCExpert compared to state-of-theart alternative methods but also its enhanced interpretability, offering a promising tool for understanding and analyzing functional brain networks.

While dFCExpert demonstrates strong performance and interpretability, several limitations offer opportunities for future improvements. First, the model has been evaluated on three large-scale fMRI datasets (HCP, ABCD, and ABIDE), but its generalizability to other datasets with different acquisition protocols, parcellation schemes, or population characteristics remains unverified. Future studies could evaluate the model on diverse datasets, including task-based fMRI, multimodal imaging (e.g., combining fMRI with EEG), or smaller datasets to assess robustness and adaptability. Second, while the modularity and state experts improve interpretability, connecting these findings to specific clinical or neurological outcomes remains a challenge. Integrating the model with clinical datasets and leveraging the learned features for diagnostic or prognostic tasks would enhance its clinical relevance. Third, the current framework relies on predefined atlases, which may not effectively capture individual variability in brain networks [50], [51]. Future research could focus on adapting dFCExpert to support personalized functional network mapping, enabling more individualized investigations of brain function. Fourth, while the model captures dFC states effectively, it does not explicitly model state transitions or the temporal relationships between states. Incorporating temporal sequence models, such as Markov chains or advanced sequence-learning architectures (e.g., transformers or RNNs), could provide deeper insights into the dynamics of state transitions. Lastly, the current implementation for sliding window adopts a window length of 50, and window stride of 3. Although it’s a standard setting for dFC analysis used in many methods [14], [26], different settings of sliding windows need to be further explored, since these settings may yield different dynamics and fluctuations for the brain activities or FCs and have different performance. Addressing these limitations through future research will enhance the robustness, efficiency, and applicability of dFCExpert, paving the way for broader adoption in both scientific and clinical settings.

## Footnotes

1 This work has been submitted to the IEEE for possible publication. Copyright may be transferred without notice, after which this version may no longer be accessible.

